# Organelle-specific lipid profiles influence/underlie metabolic health in a nutrition-dependent manner

**DOI:** 10.64898/2025.12.24.696434

**Authors:** Cassandra Tabasso, Chaitanya K. Gavini, Karin Zemski Berry, Hadi Salem, Axel Aguettaz, Sylviane Lagarrigue, Bryan C. Bergman, Virginie Mansuy-Aubert, Francesca Amati

## Abstract

Western diet (WD), characterized by high energy density, saturated fat and sucrose, is a major driver of obesity and insulin resistance (IR). Although dietary fat composition influences systemic lipid metabolism and insulin sensitivity, its impact on subcellular lipid classes distribution and fatty acid (FA) incorporation in skeletal muscle remains poorly defined. We hypothesized that (1) modulating dietary FA intake remodels mitochondrial and lipid droplet (LD) lipid profiles, including phospholipids (PL) and diacylglycerol (DAG) stereoisomers implicated in lipotoxicity; and (2) organelle-specific lipid profiles relate to metabolic health.

C57BL/6J mice were fed WD or control chow for 12 weeks. Whole-body metabolism, insulin sensitivity and substrate use were assessed by indirect calorimetry, glucose and insulin tolerance tests, and fasting biomarkers. Mitochondria and LDs were isolated from soleus muscle for organelle-resolved lipidomics. DAG isomers and PL classes were quantified, and FA chain length and saturation patterns were analyzed in total lysate (TL), mitochondria and LDs. Correlations were performed between lipid class abundance and metabolic parameters.

WD-fed mice developed obesity, dyslipidemia and early IR. Intramyocellular lipids increased, whereas mitochondrial abundance was unchanged. Organelle-resolved lipidomics revealed distinct subcellular signatures not detectable in TL. DAG FA composition mirrored dietary FA supply across compartments, with WD increasing saturated FA (SFA) and reducing di-unsaturated FA (DiUFA) species. In contrast, PL remodeling was class- and compartment-specific, with coordinated changes in mitochondria and LDs that were masked in whole-muscle TL. Several PL classes, including ether-linked phosphatidylethanolamine (ePE), phosphatidylinositol, PE and phosphatidylcholine in TL and LD, and phosphatidylserine and phosphatidylglycerol (PG) in TL and mitochondria, were strongly associated with insulin sensitivity and substrate utilization in healthy mice, but these relationships were lost under WD. LD-associated sn1,3-DAG content was a strong predictor of metabolic health in lean mice, which related to ATGL abundance. PE and PG in LD were related to obesity markers.

Dietary lipid overload induces distinct and compartment-specific remodeling of the skeletal muscle lipidome. DAG and PL classes exhibited divergent FA incorporation across mitochondria, LDs and TL, with coordinated remodeling between LDs and mitochondria that remained undetectable at the whole-muscle level. Several lipid pools, particularly LD-localized 1,3-DAG, PE and PG, were consistently related to metabolic flexibility and markers of metabolic health in healthy muscle and were disrupted in WD-fed mice. These findings identify lipid class identity, FA composition and subcellular localization as key determinants of muscle adaptation to nutritional excess and point to LD phospholipids and DAG stereoisomers as potential early molecular signatures of emerging IR.

## Introduction

Lipid metabolism is central to metabolic health, yet how subcellular lipid composition links dietary cues to insulin resistance (IR) remains unclear. Obesity and type 2 diabetes are escalating public health concerns, largely driven by Western-style diets (WD), energy-dense and enriched in saturated fat and fructose ^1,2^. These dietary patterns promote IR, through organ-specific lipid-mediated mechanisms ^3,4^. Chronic overnutrition promotes adipose tissue (AT) overload, inflammation and oxidative stress ^5^. Dysregulated AT increases lipolysis, releasing excess free fatty acids (FFAs) that accumulate ectopically. When uptake surpasses oxidative capacity, lipid intermediates including diacylglycerols (DAG) accumulate, disrupting insulin signaling via lipotoxicity ^6^, including in skeletal muscle, major determinant of whole-body insulin sensitivity (IS) ^7^. How organelles lipid profiles are affected by dietary supply, and their influence on IS remains ambiguous.

Phospholipids (PL) are the most abundant lipid family in cell membranes and are critical signaling molecules. Glycerophospholipids (GPL) consist of a glycerol backbone with two FA chains and one phosphate headgroup ^8^. Headgroup and FA characteristics modulate membrane properties, influencing organelle morphology, protein localization and stability. These features are critical for mitochondrial dynamics and LD size, which both rely on precise PL remodeling ^9,10^.

DAG, composed of a glycerol backbone and two ester-linked FA chains, exist as three stereo/regioisomers: sn-1,2-DAG, sn-2,3 DAG, and rac-1,3-DAG (further referred to as 1,2-, 1,3- and 2,3-DAG) ^11^ with different enzymatic specificity, implying distinct biological roles ^12^. In LD, TAG hydrolysis by adipose triglyceride lipase (ATGL) generates 1,3-DAG, considered biologically inactive. At the plasma membrane, 1,2-DAG arise from PL catabolism, including hydrolysis of phosphatidylinositol 4,5-biphosphate (PIP2) by phospholipases, phosphatidylcholine (PC) turnover by sphingomyelin synthase, and dephosphorylation of phosphatidic acid (PA) by lipins. *De novo* glycerolipid synthesis produces 1,2-DAG via glycerol-3-phosphate acylation and dephosphorylation ^12^. 1,2-DAG are most strongly implicated in IR through activation of novel protein kinase C (PKC) ^12,13^.

The aim of this study was to investigate how WD alters subcellular lipid composition and contributes to skeletal muscle IR. We combined metabolic phenotyping with subcellular lipidomics in mouse soleus, composed mostly of oxidative fibers and highly responsive to nutritional changes ^14^. We quantified DAG isomers and PL classes across total lysate (TL), mitochondria, and LD, and examined their associations with obesity and IR.

## Methods

### Animal care

Animal studies were conducted in accordance with the Guide for the Care and Use of Laboratory Animals of the National Institutes of Health and authorized by the veterinary authorities of the Canton of Vaud. C57BL/6J (#000664, RRID:IMSR_JAX:000664) from The Jackson Laboratory (Maine, USA) were housed in groups of four under 12:12 h light/dark cycle with *ad libitum* access to food and water. Mice temperature was monitored thrice a week. Mice were housed at 22°C with wood chip bedding and enrichment, and fed standard chow (control, Ctrl; Teklad LM-485, Envigo, Indianapolis, USA) or WD (TD88137, Teklad Diets; 42% kcal from fat, 34% sucrose by weight, and 0.2% cholesterol total) (Figure 1A) for 12 weeks ^15^. Mice were males and assigned to a group randomly at arrival (∼6weeks of age, 20±1.5gm). Experimenters were blinded during analysis according to the ARRIVE guidelines. Sample size was based on our previous studies ^15^. Two cohorts of 6 Ctrl and 6 WD were used for 1) metabolic data and lipidomics, 2) immunohistology. No adverse events were observed. Mice were euthanized through isoflurane overdose, then decapitation.

**Figure 1.**
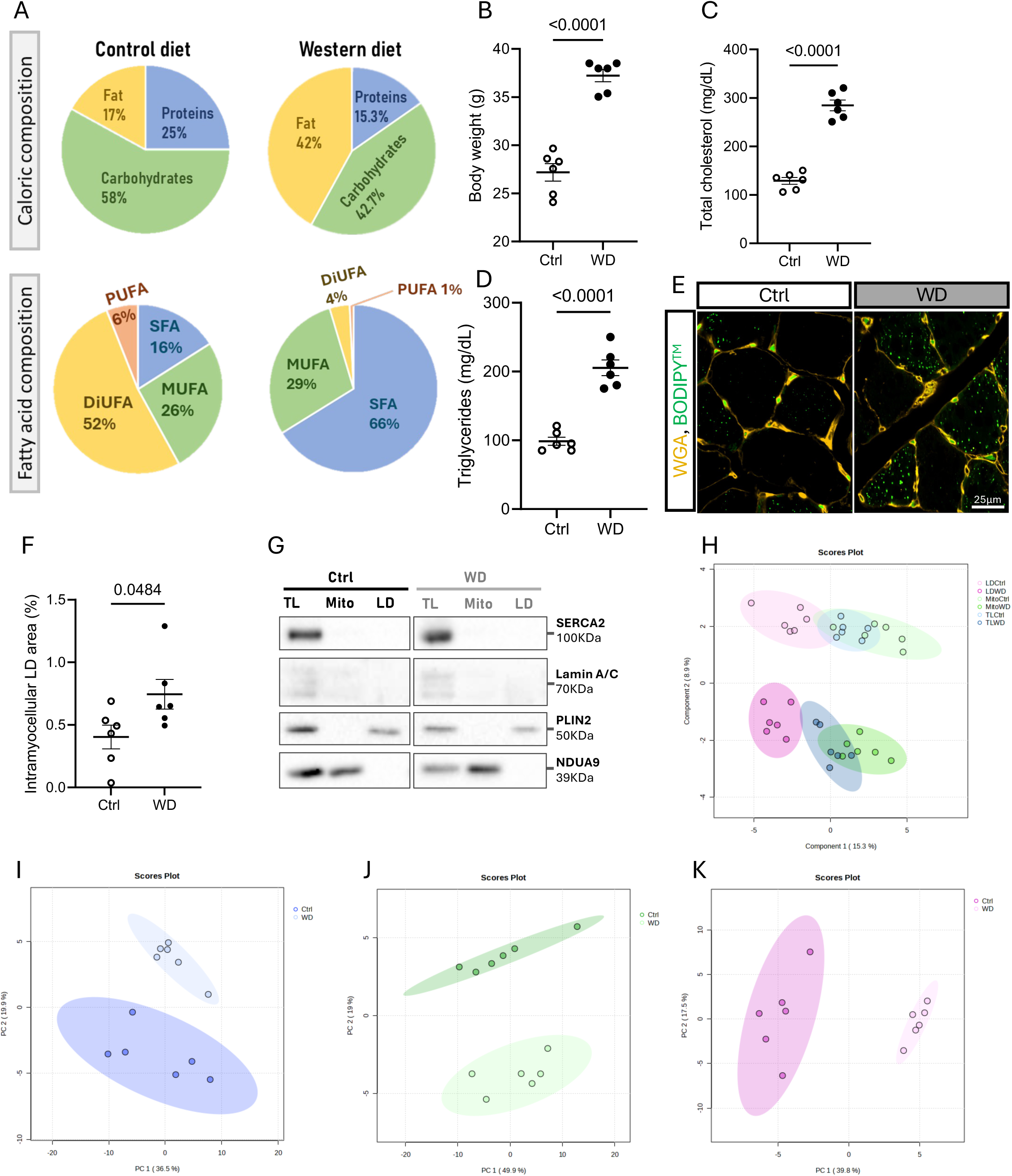
Dietary composition, metabolic phenotype, and validation of subcellular lipid fractions. (A) Macronutrients composition of control and Western diet (top) and fatty acid (FA) saturation profile within the fat fraction (bottom). (B) Body weight. (C) Plasma total cholesterol. (D) Plasma triglycerides. (E-F) LD quantification (WGA in orange, BODIPY^TM^ in green). (G) Western blotting validation of organelle isolation using compartment-specific markers: SERCA2 (endoplasmic reticulum, ER), Lamin A/C (nucleus), PLIN2 (lipid droplet, LD) and NDUA9 (mitochondria). (H) Sparse partial least squares–discriminant analysis (sPLS-DA). (I-K) Principal component analysis (PCA) of total lysate (TL, I), mitochondrial (J), and LD (K) fractions. For all panels n=6 per group, except F-G n=6 control and n=7 WD. B-D and G graphs present each sample (dots), means ± SEM. Comparisons between control (Ctrl) and WD groups were performed by two-tails independent t-test.

### Glucose and insulin tolerance tests

For glucose tolerance tests (GTT), mice were fasted overnight (12h) and administered an intraperitoneal (i.p.) glucose injection (1g/kg BW) following measurement of glucose levels. Blood glucose was monitored at 15, 30, 60, and 120min post-injection using an AlphaTrak glucometer (Fisher Scientific, Pennsylvania, USA). For insulin tolerance tests (ITT), mice were fasted for 4h and given an i.p. injection of insulin (0.5U/kg BW, Human-R Insulin U100, Lilly, Indianapolis, USA). Glucose levels were measured identically as the GTT ^16^.

### Metabolic cage analysis

Mice were individually housed in TSE PhenoMaster home-cage system (TSE systems, Chesterfield, USA) equipped for indirect calorimetry and automated feeding behavior monitoring. Temperature was maintained at 22°C using a PhenoMaster climate-controlled chamber. Experimental groups were BW-matched at the onset of metabolic assessments to minimize the confounding influence of BW on energy balance ^17^.

### Serum triglycerides, cholesterol, insulin, and leptin

Triglycerides (TAG, TR22421, Fisher Scientific), cholesterol (TR13421, Fisher Scientific), insulin, and leptin (EMD Millipore, Massachusetts, USA) were measured using manufacturer’s instructions.

### Fractionation buffer preparation

Fractionation buffer (FB) contained 20mM Tris-HCl (Biosolve Chimie, Dieuze, France), 1mM EDTA (Sigma-Aldrich, Saint-Louis, USA), and 200mM sucrose (Sigma-Aldrich), in Milli-Q water. Immediately before use, 10μL of 1000X protease inhibitor cocktail (PIC; Roche, Basel, Switzerland) and 1μL of butylated hydroxytoluene (BHT; Sigma-Aldrich; stock concentration 10mg/mL) were added per 10mL of FB. To minimize oxidative degradation, buffer was degassed by bubbling argon gas for 20sec and tubes were flushed with argon for 3sec.

### Organelle fractionation

2 Ctrl and 2 WD muscles were processed simultaneously to limit experimental bias. To prevent freeze–thaw degradation, fractions were aliquoted into duplicates for lipidomics and Western blots (WB).

Soleus (10–30mg total, both legs pooled) were homogenized in 1mL of pre-chilled FB using a Dounce homogenizer (IKA Work Janke & Kunkel, type RW 18) at 2500rpm for 25 strokes. 600μL of FB was added to homogenates.

All centrifugation steps were performed at 4°C. Lysates were centrifuged at 10,000×g for 15min. Supernatant was transferred to an ultracentrifuge tube. Pellet was resuspended in 800μL of FB and recentrifuged to recover trapped LDs; supernatant was pooled with the initial one and ultracentrifuged (Centifuge Optima L-90K, Rotor SW 60 Ti, Beckman Coulter, Brea, USA) at 100,000×g 1h, and top 2mL was LD-enriched fraction.

Pellet from the initial centrifugation was resuspended in 800μL of FB and centrifuged at 1000×g 10min. Supernatant was centrifuged twice more at 1000×g to remove debris, and resulting clarified supernatant served for mitochondrial isolation. Final pellet was resuspended in 600μL of FB, centrifuged again at 1000×g, and combined with mitochondrial supernatant. This combined fraction was then centrifuged at 10,000×g 10min to obtain the first mitochondrial pellet. Supernatant was centrifuged twice more at 10,000×g to collect additional mitochondria. All mitochondrial pellets were resuspended in 100μL of FB and pooled for final mitochondrial fraction.

### Fractions validation

WB were performed as previously described ^18^ on 5 sets of biological replicates per group. 5μg of protein from TL, mitochondrial and LD fractions were loaded. Memcode^TM^ Total Protein Stain (Sigma-Aldrich) was used for loading control. Primary antibodies validated by suppliers were incubated overnight at 4°C: rabbit anti Lamin A/C (1:1000, RRID:AB_648154, sc-20681 (H-110), Santa Cruz), rabbit anti-SERCA2 (1:1000, RRID:AB_2758348, A1097, Abclonal, Woburn, USA) rabbit anti-PLIN2 (1:1000, RRID:AB_10863476, ab108323 (EPR3713), Abcam, Cambridge, UK), and mouse anti-NDUA9 (1:2000, RRID:AB_301431, ab14713, Abcam). Primary antibodies were diluted in 2% Bovine serum albumin (BSA, Sigma-Aldrich), except anti-Lamin A/C, diluted in 5% skim milk. HRP-conjugated anti-mouse IgG (1:5000, RRID:AB_330924, 7076, Cell signaling) and anti-rabbit IgG (1:5000, RRID:AB_2099233, 7074, Cell signaling) were incubated 2h at RT. Signal acquisition was performed with Biorad Chemidoc (ChemiDoc XRS+, Bio-Rad Laboratories). Due to limited biological material, WB was performed once.

### ATGL quantification

5μg of proteins from 6 LD samples per group were loaded. Anti-ATGL (1:1000, RRID:AB_2167955, 2138, Cell signaling) was added, followed by anti-rabbit IgG. Band intensity was normalized to Memcode^TM^ and quantified using Fiji (National Institutes of Health) ^19^.

### Lipid extraction and lipidomics

Samples were processed randomly by fraction to avoid bias. After addition of internal standard cocktail, TL, LD, and mitochondrial lipids were extracted with methanol and methyl tert-butyl ether ^20^. TAG, DAG, sphingolipids, and PL were analyzed as described previously ^21,22^. Concentrations were determined using stable isotope dilution with standard curves for saturated and unsaturated lipids.

### Lipid data analyses

Lipid concentration was normalized to tissue wet weight. Individual species were summed for class abundance. To assess FA composition, lipid concentrations were duplicated to evaluate FA chains individually. Lipid species were classified by saturation as saturated FA (SFA), monounsaturated FA (MUFA), di-unsaturated FA (DiUFA) and polyunsaturated FA (PUFA), and by length. Relative proportions were computed.

### LD and mitochondria quantification

1 Ctrl and 1 WD sample were processed simultaneously. Soleus were fixed in 4% formaldehyde (Carl Roth, Karlsruhe, Germany) and PBS at 20-25°C 2h, then at 4°C 16h. After embedding in Cryomatrix^TM^ (Epredia, Tokyo, Japan), samples were frozen in cooled isopentane (Sigma-Aldrich) 30min, then immersed in liquid nitrogen and stored at -70°C.

Soleus were processed as previously described ^23^. Anti-TOMM20 was diluted in PBS-BSA 2% and incubated 16h at 4°C (1:1000, RRID:AB_2207530, 11802-1-AP, Proteintech, Rosemont, UK). Donkey anti-Rabbit IgG (H+L) Alexa Fluor™ 647 (1:500, RRID:AB_2536183, A-31573, Thermo Scientific) was incubated 4h at 20-25°C. Wheat Germ Agglutinin Texas Red (WGA) (1:200, W21405, Thermo Scientific) and BODIPY™ 493/503 (1:500, D3922, Invitrogen, Massachusetts, USA) in PBS were applied 1.5h at 20-25°C. Hoechst (1:8000, H3570, Invitrogen) was added in PBS 3min. Slides were washed thrice in PBS between steps. Sections were mounted with Mowiol on 75×25mm glass slides (Epredia) and covered with 24×60mm coverglass (Merck, Darmstadt, Germany).

Images were acquired blinded using a spinning disk confocal microscope (Nikon Ti2, CrEST Optics aX-Light V3, Nikon, Tokyo, Japan) with 60X objective in immersion oil (CFI Plan Apochromat Lambda 60X Oil, N.A. 1.40, W.D. 0.13mm, Nikon).

Image analyses were performed blinded. 2 images per mouse were averaged. In Fiji ^19^, regions of interest (ROI) around myofibers were drawn using WGA and combined into a single ROI. Intracellular-extracellular areas were separated. BODIPY™ staining was manually thresholded to select lipid depositions. Using the “analyze particles” function, BODIPY™ was quantified for each mask separately. BODIPY™ staining was normalized by mask area: 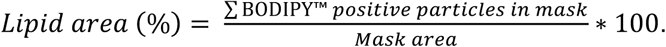 For mitochondria, TOMM20 positive area was thresholded and normalized by intracellular mask area.

### Statistical analyses

Data are means±SEM. After assessing normality and homogeneity of variance, group comparisons two-tailed independent *t*-tests were performed. If data was not normal, Mann-Whitney tests were performed. Organelle fractions were analyzed independently. For FA chain length and saturation comparisons within lipid classes, two-way ANOVA was performed with diet as the main effect, followed by Šídák’s multiple comparisons test. Species analyses were performed with multiple t-tests, with 1% FDR and two-stage step-up method of Benjamini, Krieger and Yekutieli. Sample sizes refer to biological replicates. P-value<0.05 was considered statistically significant. Unless otherwise indicated, statistical analyses and graphs were performed using GraphPad Prism9 (GraphPad, La Jolla, USA).

Multivariate analyses of lipidomics data were conducted using principal component analysis (PCA) and sparse partial least squares–discriminant analysis (sPLS-DA) with *MetaboAnalys*t ^24^. PCA, an unsupervised dimensionality reduction method, was used to summarize overall lipid variance and sample clustering by generating principal components (PCs), capturing major variation axes. PC1 and PC2 are first and second components, accounting for the largest and second-largest proportions of total variance. SPLS-DA, a supervised approach, identified specific lipid variables that discriminate between predefined groups (i.e. diet, organelle fraction). “Sparse” feature selection step limits the number of variables included, highlighting the greatest contribution to group separation and minimizing overfitting. These complementary analyses were used to provide both unsupervised (PCA) and supervised (sPLS-DA) insights into WD effects on lipid composition. Settings were as follows: missing values were imputed with one-fifth of minimum detected value per variable, no filtering for reliability, variance and abundance, data were log-transformed and Pareto-scaled, 95% confidence intervals were displayed as colored ellipses. LION/web ^25^ was used for lipid enrichment. Ctrl vs WD and WD vs Ctrl were compared, and FDR q-value <0.05 was considered significant.

## Results

### WD induces metabolic syndrome-like features and alters subcellular lipid composition in skeletal muscle

FA profile of WD contained a greater SFA proportion and lower DiUFA and PUFA (Figure 1A). WD-fed mice were heavier (Figure 1B), had elevated circulating cholesterol and TAG (Figure 1C-D). Intramyocellular lipids increased (Figure 1E-F), without changes in extracellular lipids or mitochondria (Figure S1A-C).

After WB validation (Figure 1G), organelle fraction drove separation along component 1 (15.3%) through sPLS-DA, whereas component 2 (8.9%) showed diet-induced variations (Figure 1H). In PCA of TL, mitochondria, and LD, PC1 explained 36.5%, 49.9%, and 28.9% of the variance, respectively (Figure 1I–K), with PC2 reflecting further dietary separation.

### Obesity- and IR-related traits correlate with subcellular lipid profiles in a diet-dependent manner

WD-fed mice displayed higher fat-to-lean ratio (Figure S1D) and increased fat and lean mass (Figure S1E-F), indicative of diet-induced obesity. Circulating leptin concentrations increased in WD-fed mice (Figure S1G), indicating leptin resistance.

GTT and ITT revealed no differences in dynamic responses (Figure 2A-B, Figure S1H-I). WD displayed higher fasting glucose and insulin (Figure 2C, Figure S1J), consistent with early-stage IR, potentially preceding overt muscle IR.

**Figure 2.**
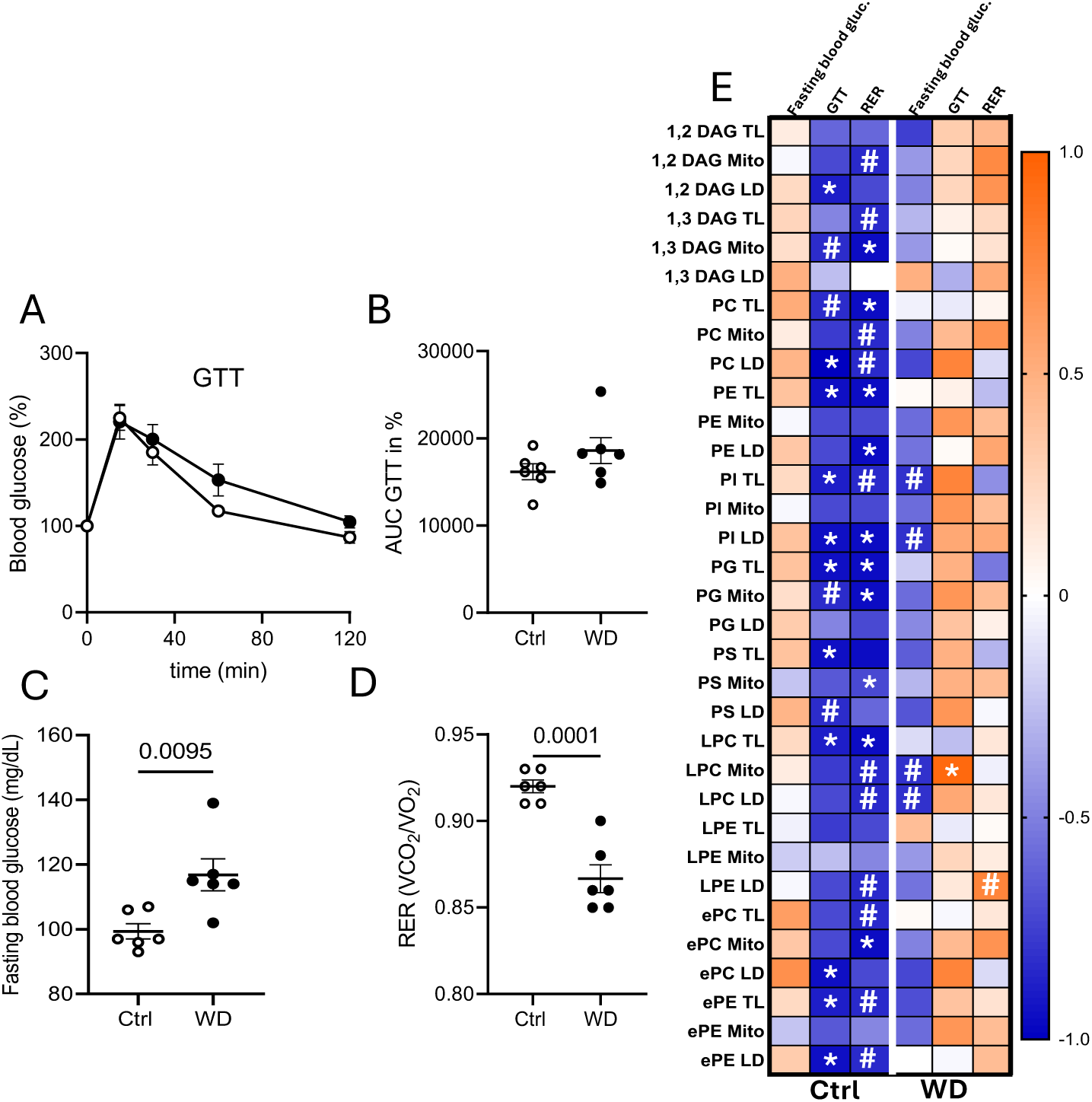
Relationships between metabolic traits and subcellular lipid classes. (A-D) Metabolic parameters including GTT glucose profile (A), GTT area under the curve (B), fasting blood glucose (C), and respiratory exchange ratio (RER)(D). n=6 per group. A, C and D graphs present each sample (dots), means ± SEM. Comparisons between control (Ctrl) and WD groups were performed by independent t-test. (E) Spearman correlation heatmap of lipid class abundance with metabolic parameters. Blue indicates negative correlations and orange indicates positive correlations. * p-value <0.05, # p=0.05-0.1. For all panels n=6 per group.

Respiratory exchange ratio (RER) revealed a diet-dependent shift in substrate utilization. Ctrl averaged a RER of 0.92, reflecting predominant carbohydrate oxidation, while WD averaged 0.86, indicating increased FA reliance ^26^(Figure 2D).

Correlation analyses revealed distinct associations between lipid composition and metabolic traits (Figure 2E). In Ctrl TL, several PL classes negatively correlated with GTT, including phosphatidylethanolamine (PE), phosphatidylinositol (PI), phosphatidylglycerol (PG), phosphatidylserine (PS), lyso-phosphatidylcholine (LPC) and ether-linked PE (ePE), suggesting better glucose tolerance with higher PL abundance. Similarly in LD, GTT values were inversely associated with 1,2-DAG, PC, PI, ePC, and ePE. LD ceramides were borderline to positive correlations with fasting insulin and ITT (Figure S2A).

In WD, associations were largely lost. Isolated and borderline significant correlations persisted, including negative relationships between fasting blood glucose and PI in TL and LD, and LPC in mitochondria and LD (Figure 2E). LD 1,3-DAG correlated with ITT while mitochondrial LPC positively correlated with GTT (Figure 2E, Figure S2B), consistent with reported pro-inflammatory effects ^27^.

In Ctrl, fasting RER negatively correlated with several PL, including PC, PE, PG, and LPC in TL; 1,3-DAG, PG, PS, and ether-linked PC (ePC) in mitochondria; and PE and PI in LD (Figure 2E), indicating that low abundance associated with higher RER. Mitochondrial ceramides negatively correlated with fat mass in healthy mice (Figure S2C).

In contrast, WD showed no consistent relationships between RER and lipid abundance (Figure 2E). Isolated associations persisted, such as PI in TL negatively correlated with plasma cholesterol, and mitochondrial LPC with fat mass (Figure S2D). In LD, PC and ePC inversely correlated with BW (Figure S2B).

### PE and PG in LD are biomarkers of metabolic health independently of diet

To assess whether some relationships between lipids and metabolism were diet independent, we pooled all mice (Figure 3A-B). Certain lipid species reflected IS status: LD PE correlated positively with fasting insulin. LD PE appeared as robust obesity markers through positive associations with BW, plasma triglycerides, and leptin, and negative with RER (Figure 2G). LD PG correlated with fasting insulin. Both PE (Figure 3C) and PG (Figure 3D) increased in WD LD fractions.

**Figure 3.**
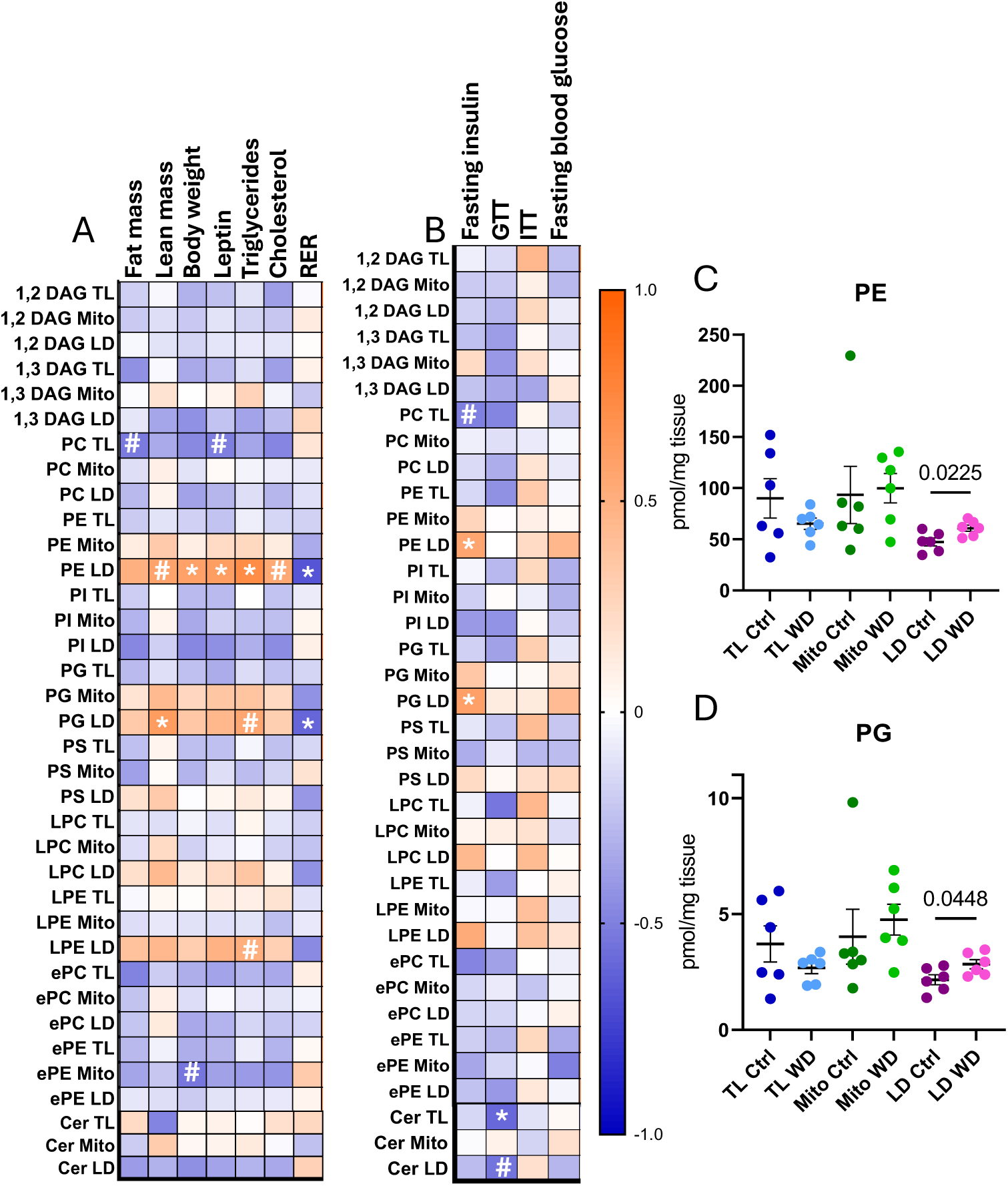
**Insulin sensitivity and body type correlates to PE and PG in LD**. (A-B) Spearman correlations between lipid abundance in TL, mitochondrial and LD fractions and obesity indicators (A) or insulin sensitivity markers (B). Blue indicates a negative correlation and orange indicates a positive correlation. * p-value < 0.05 and # p= 0.05-0.1. (C-D) Relative abundance across factions of PE (C) and PG (D). C- D graphs present each sample (dots), means ± SEM. Comparisons between control (Ctrl) and WD groups were performed by independent t-test. For all panels n=6 per group.

### 1,3-DAG in LD are indicators of metabolic health

LD-associated 1,3-DAG inversely correlated with BW, plasma leptin and cholesterol in healthy mice (Figure 4A-B), pointing to lean metabolic phenotype. We assessed ATGL abundance in LD fractions (Figure 4C) and found no significant differences (Figure 4D). In Ctrl, ATGL positively correlated with LD 1,3-DAG. In WD, ATGL and 1,3-DAG levels correlated negatively. ATGL abundance did not correlate to 1,2-DAG in any fraction, indicating selective associations.

**Figure 4.**
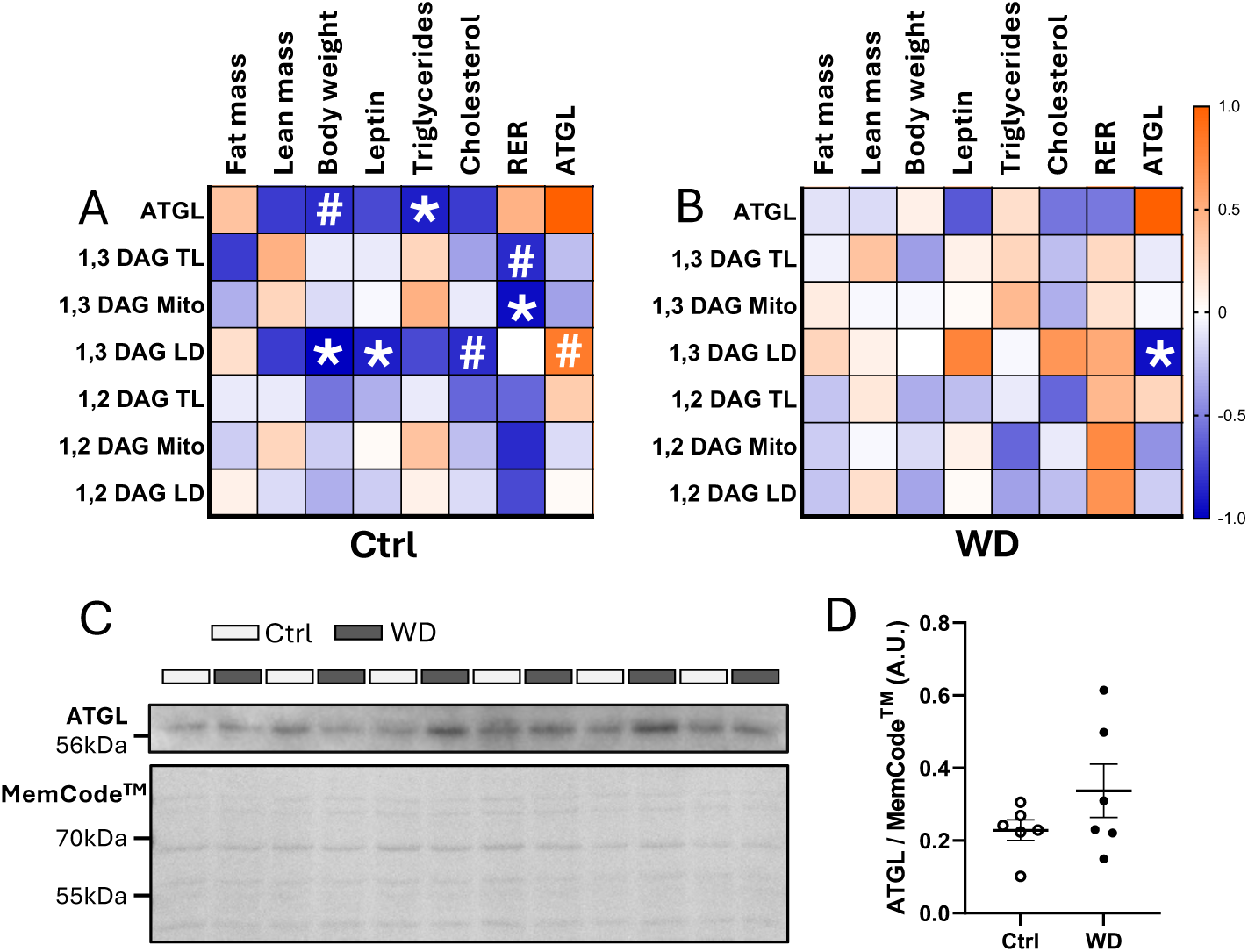
1,3-DAG correlates with metabolic status and ATGL expression in a diet-specific manner. (A-B) Spearman correlations between 1,2- and 1,3-DAG and obesity-related parameters in (A) control and (B) WD-fed mice. Blue indicates negative correlations and orange indicates positive correlations. * p-value < 0.05 and # p=0.05-0.1. (C) Western blot analyses of ATGL in LD fractions. (D) Quantification of ATGL abundance. Comparison between diets were performed by two-tailed t-test. For all panels n=6 per group.

### Diet modulates FA remodeling of DAG species

1,2-DAG species containing 14:0 FA consistently increased in TL and LD of WD (Figure 5A-B). LION enrichment analysis (Table S1) suggests that 14:0 FA, SFA and MUFA drive WD phenotype, while 18:2 FA and PUFA drive the healthy phenotype in Ctrl. We subsequently assessed FA chains composition to identify diet-specific patterns. FA chain remodeling was consistent across subcellular fractions (Figure 5C, Table 1, Supp. Table 1). WD increased 16C FA and reduced 18C FA in TL, mitochondria and LD, with increase in 14C FA in LD. SFA and MUFA were elevated with WD, while DiUFA decreased.

**Figure 5.**
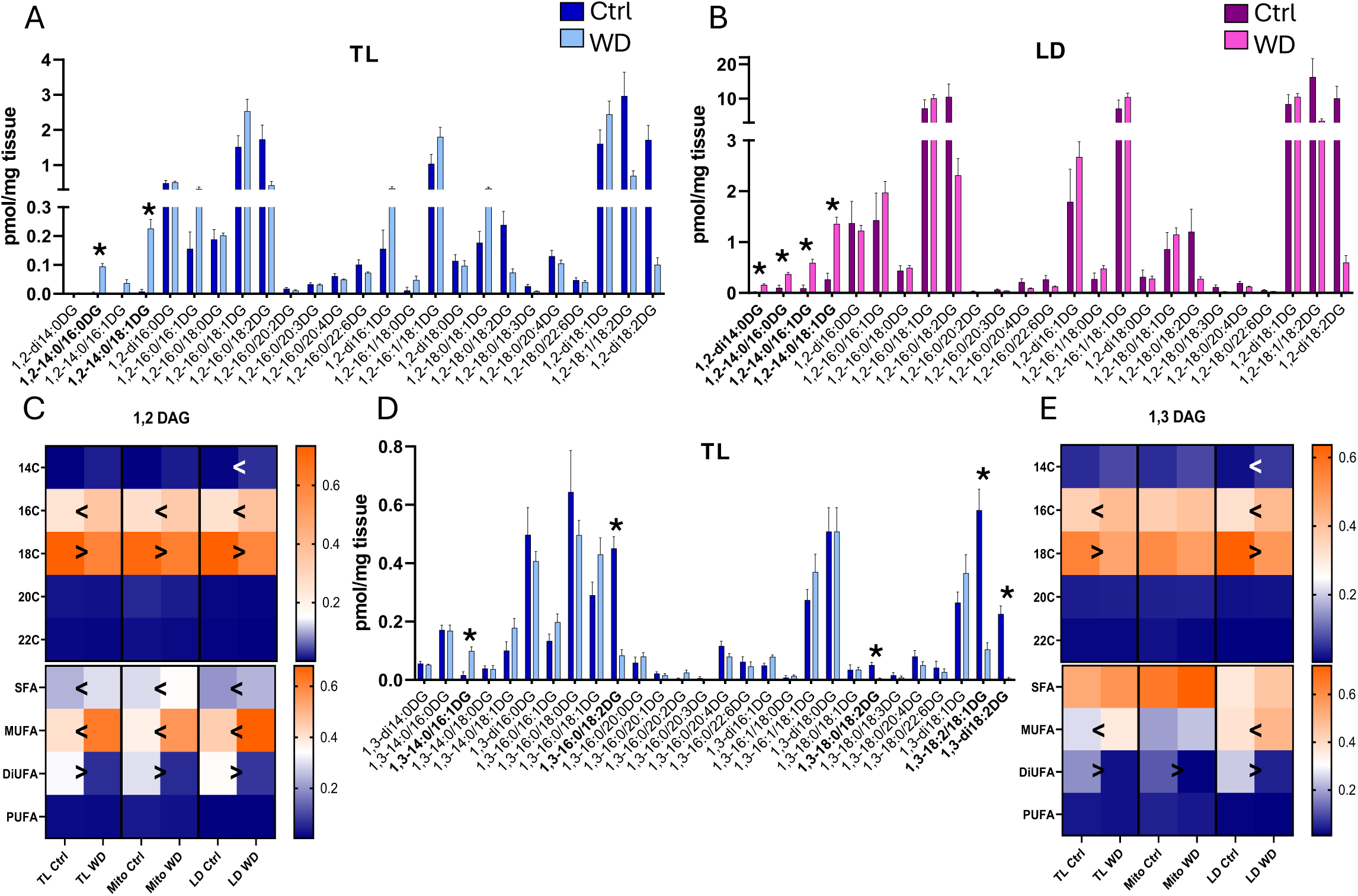
DAG species and FA chains composition. (A-B) 1,2-DAG species in (A) TL and (B) LD. (C) 1,2-DAG FA chain length (top half of each panel) and saturation (bottom half of each panel). (D) 1,3-DAG species in TL. (E) 1,3-DAG FA chain length (top half of each panel) and saturation (bottom half of each panel) for 1,2-DAG. (A,B,D) Comparisons between groups were performed through multiple t-tests. *p<0.05. (C,E) Comparisons between diets were performed by two-way ANOVA with each fraction analyzed separately. n=6 per group. Blue indicates proportions close to 0 and orange proportions close to 1. < and > symbols denote p-values<0.05, with the arrow showing the direction of the change. For all panels n=6 per group.

**Table 1.**
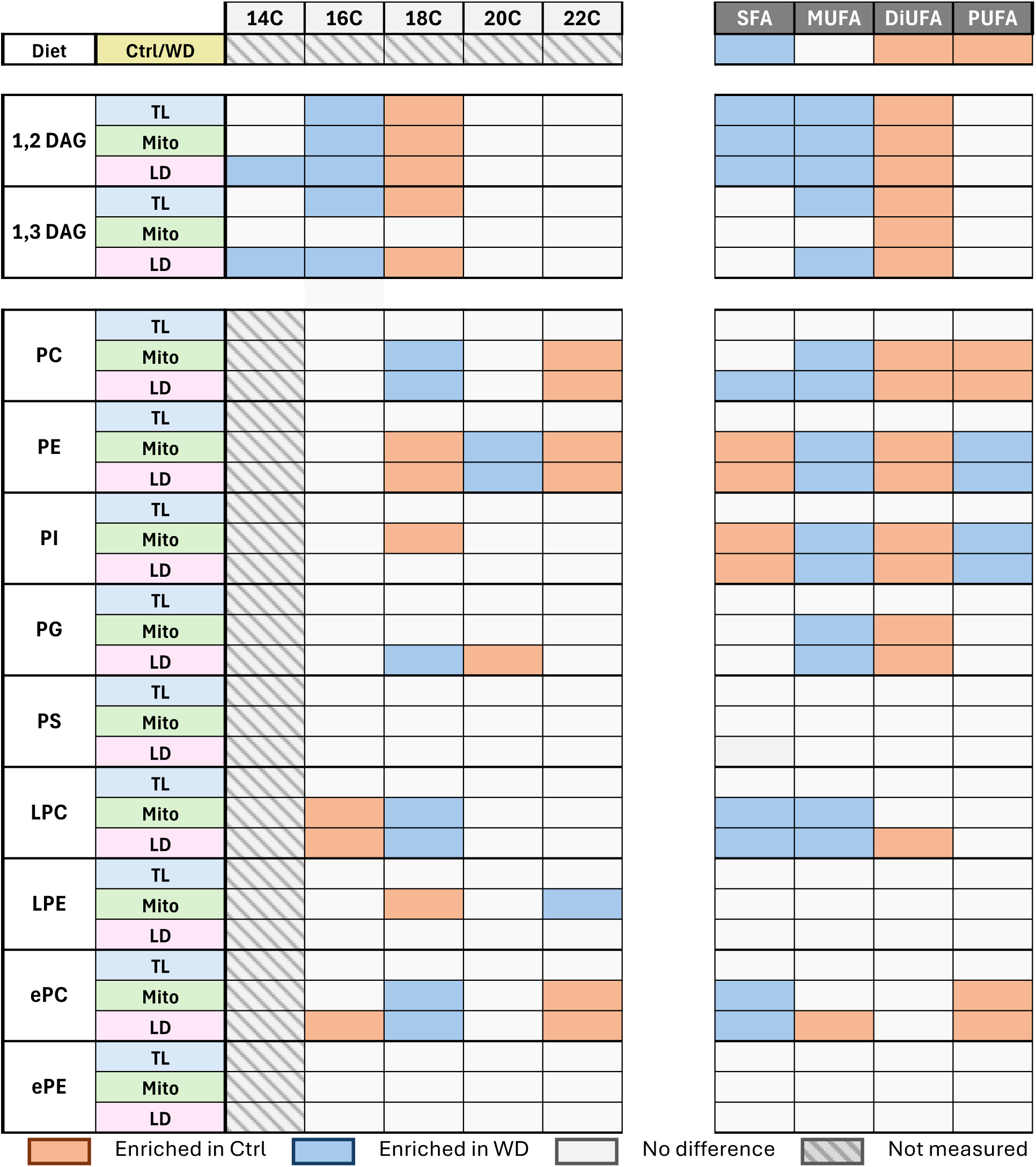
Summary of diet-modulated fatty acid chain integration. . Notes: In colors, significant dilerences with p- value <0.05. In gray, p- value >0.05. In patterns, FA lengths not assessed during lipidomics. Abbreviations: TL total lysate, LD lipid droplet, Mito mitochondria, FA fatty acid, SFA saturated fatty acid, MUFA monounsaturated fatty acid, DiUFA diunsaturated fatty acid, PUFA polyunsaturated fatty acid, Ctrl control, WD western diet.

1,3-DAG species containing 14:0 FA increased in TL of WD (Figure 5D), while 1,3-DAG containing 18:2 FA increased in Ctrl. No changes were observed in LD or mitochondria (data not shown). FA remodeling was similar in TL and LD but not in mitochondria (Figure 5E, Table 1). WD increased 16C FA and MUFA, while reducing 18C FA and DiUFA, mirroring dietary composition. In mitochondria, Ctrl showed increased DiUFA, with no other FA changes.

### PL remodeling is class- and compartment-specific

PL species changed with diet only in LD and species-specific patterns (Figure 6A-G). In WD, we observed increases of FA16:/18:0 in PC, PE and PG, 18:0/18:1 in PC, ePC, PE, PI and PG, di18:1 and 18:1/20:4 in PC, PE, PI and PG, and 18:1/22:5 in PC, PE and PG. Several other combinations of FA containing at least one SFA or MUFA increased in WD LD, including 18:1/22:6 in PS and 18:1 in LPC. In Ctrl, FA combinations containing 18:2, like di18:2 in PC and PE, and 18:0/18:2 in PC and PI consistently increased.

**Figure 6.**
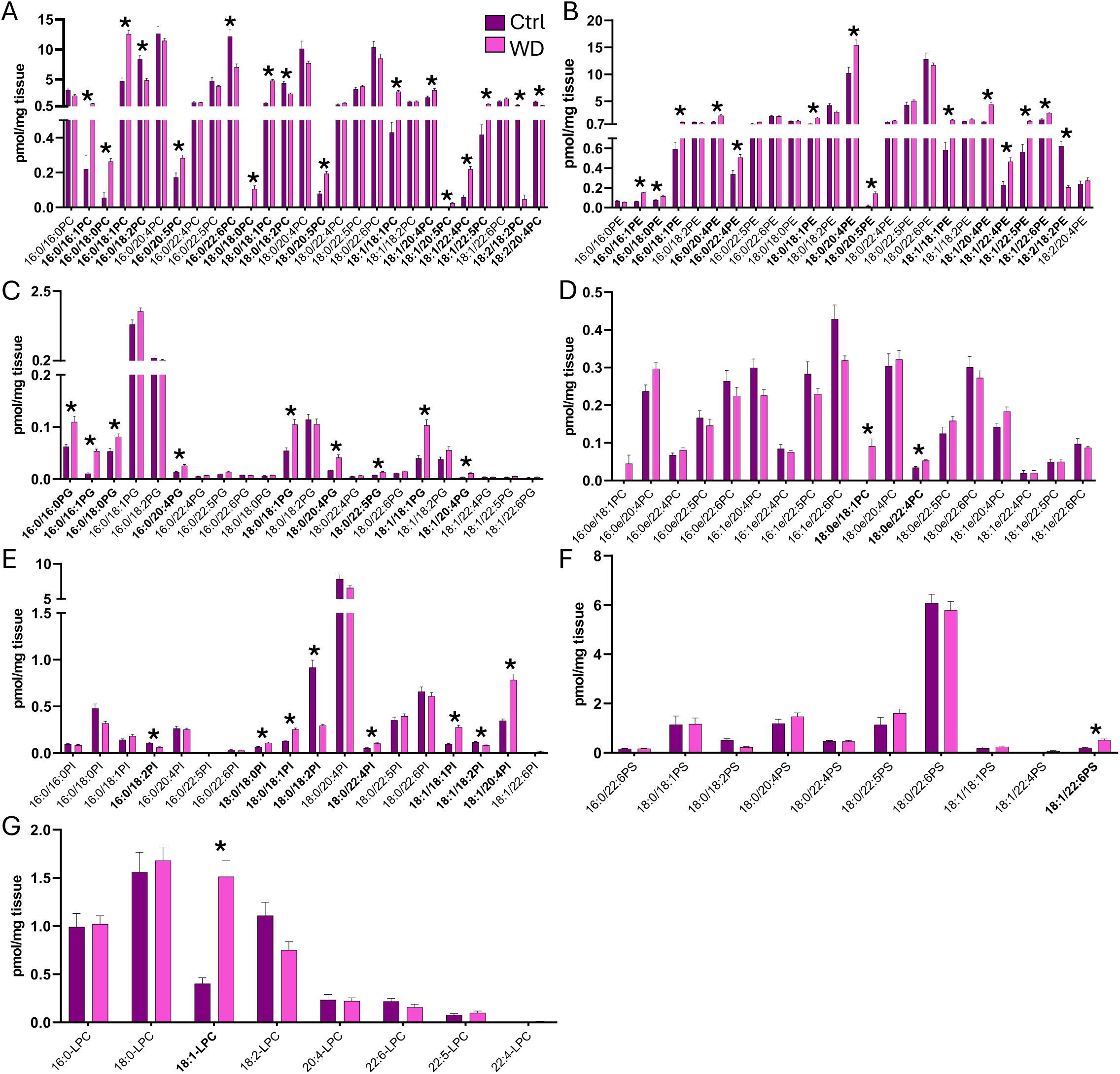
LD PL species. (A-G) Species in (A) PC, (B) PE, (C) PG, (D) ePC, (E) PI, (F) PS and (G) LPC. Comparisons between groups were performed through multiple t-tests. *p<0.05.

PL FA remodeling was compartment- and class-dependent, with mitochondria and LD showing similar trends (Figure 7A-G, Table 1). 16C FA decreased in ePC LD and LPC LD and mitochondria of WD. 18C FA increased in WD in mitochondrial and LD ePC, LPC, PC and in LD PG, but decreased in mitochondrial PI and LPE, and LD and mitochondrial PE. WD also increased 20C FA in mitochondrial and LD PE, and decreased in LD PG. Finally, 22C FA decreased in WD in mitochondrial and LD ePC, PC and PE, but increased in mitochondrial LPE.

**Figure 7.**
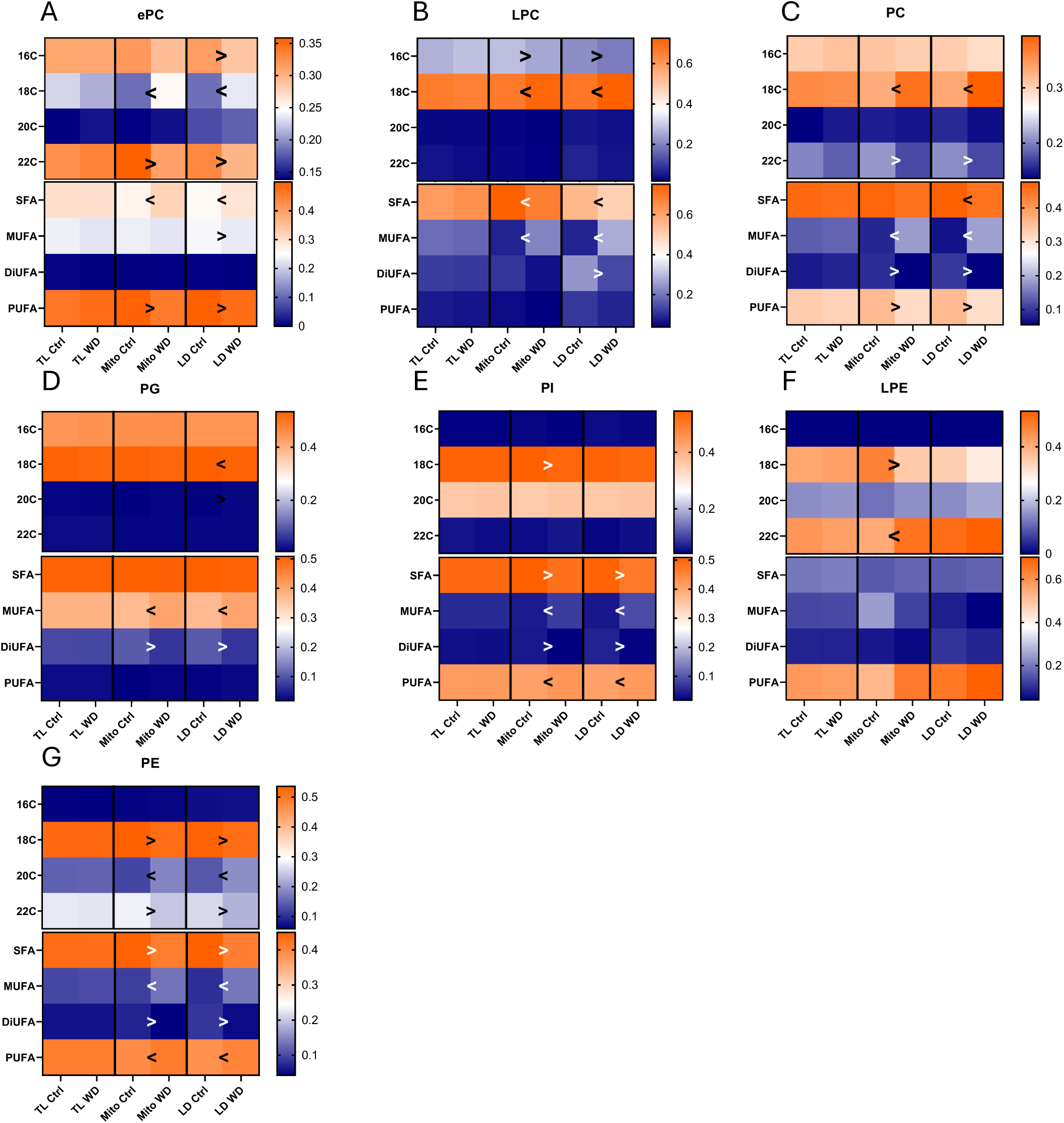
FA chains composition of PL classes across subcellular fractions. (A-G) FA chain length (top half of each panel) and saturation (bottom half of each panel) for individual PL classes. Comparisons between diets were performed by two-way ANOVA with each fraction analyzed separately. n=6 per group. Blue indicates proportions close to 0 and orange proportions close to 1. < and > symbols denote p-values<0.05, with the arrow showing the direction of the change. For all panels n=6 per group.

SFA increased in WD in LD PC (Figure 7C), and mitochondrial and LD ePC and LPC (Figure 7A-B). Unexpectedly, Ctrl displayed higher mitochondrial and LD SFA in PI and PE (Figure 7E,G), contrary to diet. MUFA increased in WD in mitochondrial and LD LPC, PC, PG, PI and PE (Figure 7B-E,G), but decreased in LD ePC (Figure 7A). DiUFA decreased in WD in mitochondria and LD in ePC, PC, PG, PI, PE (Figure 7A,C,E,G), and in LD LPC (Figure 7B), while increasing in Ctrl, consistent with dietary composition. PUFA decreased in WD in mitochondrial and LD ePC and PC (Figure 7A,C), but increased in PI and PE (Figure 7D,G).

## Discussion

WD is widely used in murine models because it reliably induces metabolic disturbances and closely mimics modern human dietary patterns, particularly regarding lipid overload, insulin signaling defects and fat accumulation. While lipidomic analyses characterized WD-induced hepatic steatosis and systemic dyslipidemia ^28–30^, comprehensive assessment of muscle-specific lipid remodeling and its relationship to metabolic phenotypes has been limited. To address this gap, we combined in vivo metabolic phenotyping with organelle-resolved lipidomics to determine how dietary lipid supply shapes subcellular lipid composition in skeletal muscle. This approach allowed us to dissect lipid remodeling within mitochondria, LDs and TL, revealing distinct compartment-specific signatures, hidden in whole tissue. We identified lipid classes in LD (e.g. PE, PG) and DAG stereoisomers (1,3-DAG) that consistently associated with IS, substrate utilization and body composition in a diet-dependent manner. Our findings demonstrate that DAG fatty acid composition follows dietary supply across compartments, while PL remodeling is class- and compartment-specific and therefore masked in whole-tissue analyses, underscoring the need for subcellular resolution. Identifying lipid classes and organelle-specific lipid pools linked to early metabolic impairment may support the development of lipid-based biomarkers or therapeutic targets for the prevention or early detection of IR.

PL classes including PC, PE, PI, PG, PS, LPC, ePC and ePE emerged as correlates of IS and body composition in healthy mice, showing inverse associations with glucose intolerance and RER. Because lower RER reflects greater fatty acid oxidation (FAO), these correlations suggest that higher PL abundance, particularly within mitochondria and LD, is related to enhanced FAO and metabolic flexibility in lean, healthy animals. Several of these PL classes play established roles in mitochondrial function: PG contributes to cardiolipin synthesis and FAO, PS serves as precursor for PC and PE synthesis, PE regulates cristae architecture and mitochondrial biogenesis, and ePC supports redox balance ^10,31^. Taken together, these PL species serve as indicators of mitochondrial integrity and oxidative capacity.

Lipid composition also strongly influences LD function, where both the neutral lipid core and PL monolayer determine droplet size, protein recruitment, and organelle contact dynamics. PE- and DAG-enriched LDs recruit diacylglycerol O-acyltransferase 2 (DGAT2), promoting de novo lipogenesis. PE-rich LDs also attract anti-lipolytic proteins and form large LDs typical of muscle lipid infiltration (myosteatosis) observed in obesity ^32,33^ and reproduced here in WD mice. This reinforces PE as structural marker of obesity-related LD remodeling ^34,35^, and its potential association to IR. In our study, LD PG correlated positively with lean mass, which was higher in WD-fed mice and associated with lower RER, consistent with observations in human obesity ^36^. LD fractions may include membrane contact sites, which does not diminish the physiological relevance of the associations, as PG presence likely reflects increased mitochondrial-LD interactions, which are observed under metabolic stress and elevated FAO ^37^.

1,3-DAG abundance in LDs emerged as a prominent feature of body morphotype traits. Although 1,3-DAG species are often deemed biologically inactive, because they cannot activate PKCs and have shown limited correlation with IS in previous reports ^38,39^, our findings indicate a potential protective role. This interpretation aligns with a nutritional study showing that dietary supplementation with DAG oil attenuated WD-induced IR and enhances transcription of FAO-related genes in skeletal muscle ^40^. 1,3-DAG accumulation in LDs of healthy mice may therefore indicate efficient TAG turnover by ATGL coupled to FAO, shown to relate to IS through decreased ability to produce lipotoxic intermediates ^41^. ATGL expression is elevated in oxidative fibers and endurance-trained muscle ^13,42^, where it supports mitochondrial oxidative capacity. The inverse association between 1,3-DAG and obesity markers in Ctrl therefore suggests that LD 1,3-DAG reflect metabolic efficiency and active lipid turnover.

for FAO, particularly in oxidative fibers like soleus. Conversely, the loss or reversal of ATGL-1,3-DAG correlations in WD-fed mice may indicate impaired TAG-DAG cycling, enhanced re-esterification, or decreased TAG synthesis, underscoring disrupted lipid flux in IR and obesity. Interestingly, 1,2-DAG, typically considered as lipotoxic, showed no association with metabolic parameters in either healthy or insulin-resistant mice. Similarly, ceramides, despite being classical mediators of mitochondrial dysfunction, correlated negatively with fat mass only in Ctrl, suggesting context-dependent effects that may become more relevant in advanced metabolic disease. In WD-fed mice, LD ceramides trended positively with IR markers, consistent with a transitional metabolic state (IR without overt diabetes). Muscle type and subcellular localization are critical determinants of lipid-metabolism interactions. Soleus contains approximately 80% oxidative, slow-twitch fibers, which inherently store more lipid than glycolytic muscles. As a result, our findings may be specific to oxidative tissue and may not capture the same “lipotoxic candidates” typically identified in more glycolytic muscle types.

Further than class abundance, FA chain composition also modulates organelle morphology and function. In our dataset, FA composition of 1,2-DAG was conserved across TL, mitochondria and LD, and across TL and LD for 1,3-DAG, suggesting that its remodeling is primarily diet-driven. Specific candidates emerged as drivers for that change, notably 14:0 FA, which has been associated to IR and obesity in human muscle^43^. In contrast, PL FA composition was compartment-specific, with coordinated remodeling in mitochondria and LD but minimal changes at the whole-cell level, pointing to shared regulatory mechanisms like PL or FA trafficking through organelle contacts. In Ctrl LD, 18:2 FA consistently increased, reflecting association of 18:2 FA to healthy diet and exercise training in rat muscle ^44^. In WD, LD PL species had higher combinations of 16:0, 16:1, 18:0 and 18:1 FA, suggesting possible modulation by Stearoyl-CoA Desaturase 1 (SCD1) and Elongase 6 (ELOVL6), both associated to obesity induced IR ^45–47^. DAG and PL species from WD-fed mice were enriched in SFA and MUFA. Saturated PL tend to enlarge LDs by decreasing monolayer fluidity, a phenomenon demonstrated primarily in *in-vitro* models ^35^ but that is consistent with the increased LD area in WD mice. Within mitochondria, FA composition influences mitochondrial function through microdomain formation and modulation of electron transport chain efficiency ^48–50^. These observations highlight the importance of assessing lipid saturation at the subcellular level to better understand organelle adaptation in metabolic disease.

In summary, our findings demonstrate that the relationship between lipid composition and metabolic phenotype is nonlinear and context dependent. This relationship is robust in metabolically healthy muscle but becomes disrupted under obesity and IR. This study underscores the importance of lipid class identity, FA composition, and subcellular localization in determining metabolic outcomes and highlights how dysregulated lipid remodeling contributes to diet-induced metabolic dysfunction.

## Authorship

C.T. researched data, contributed to the study concept, design, analyses, visualization and wrote the draft of the article. C.G., K.Z.B, H.S., A.A. and S.L. researched the data and edited the article. B.B. and V.M.A contributed to the study concept, interpretation of the data and edited the article. F. A. contributed to the study concept, design, analyses, interpretation of the data; and wrote the article. B.B., V.M.A and F.A. supported funding for all personnel and methods of this study. All authors reviewed and approved the final manuscript.

## Acknowledgments

We thank Julien Duvernay for his technical assistance and the Cell Imaging Facility of the Faculty of Biology and Medicine, University of Lausanne. This study was supported by the Swiss National Science Foundation (grants: 320030_170062 and 310030_188789, IZCOZ0_205425, all to FA) and the Colorado Nutrition Obesity Research Center grant (P30DK048520 to BB).

## Data availability

All data supporting the findings of this study are available within the paper and its Supplementary Information. Lipidomics data are available from the corresponding author upon reasonable request.

---

**Supplemental Figure 1.**
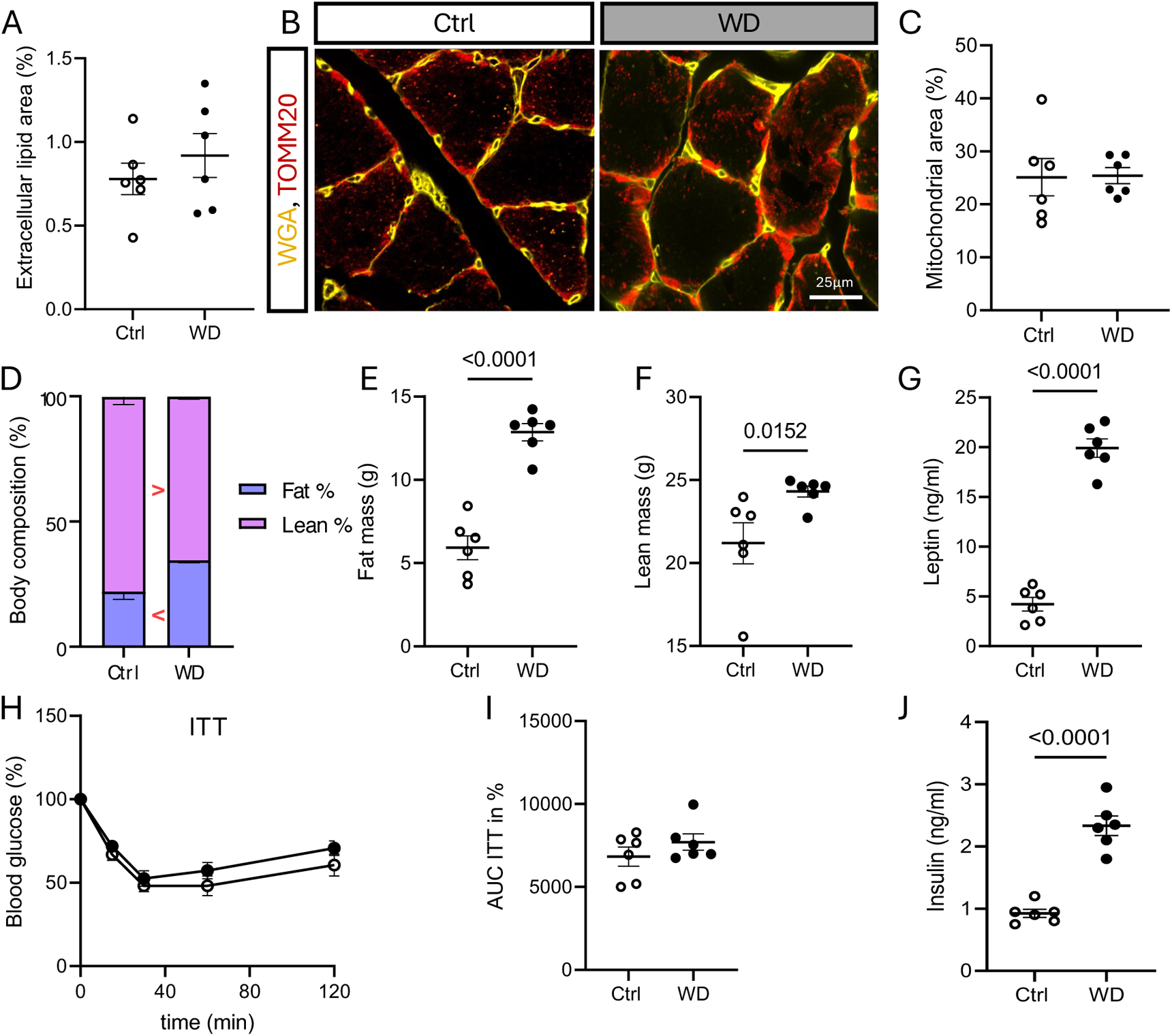
Body composition and metabolic parameters in WD-fed and control mice: (A) Extramyocellular lipids quantification. (B-C) Mitochondrial quantification (WGA in yellow, TOMM20 in red) (B) and mitochondrial area (C). (D-F) Body composition analysis showing fat-to-lean ratio (D), absolute fat mass (E), and lean mass (F). (G) Plasma leptin concentrations. (H-I) Insulin tolerance test (ITT) and corresponding area under the curve (AUC). (J) Plasma insulin levels. For panels A-H n=6 per group. For panels I-J n=6 control and n=7 WD. All graphs present each sample (dots), means ± SEM. Comparisons between control (Ctrl) and WD groups were performed by independent t-test. For panel A: > and < symbols indicate the direction of group dilerences (p < 0.05).

**Supplemental Figure 2.**
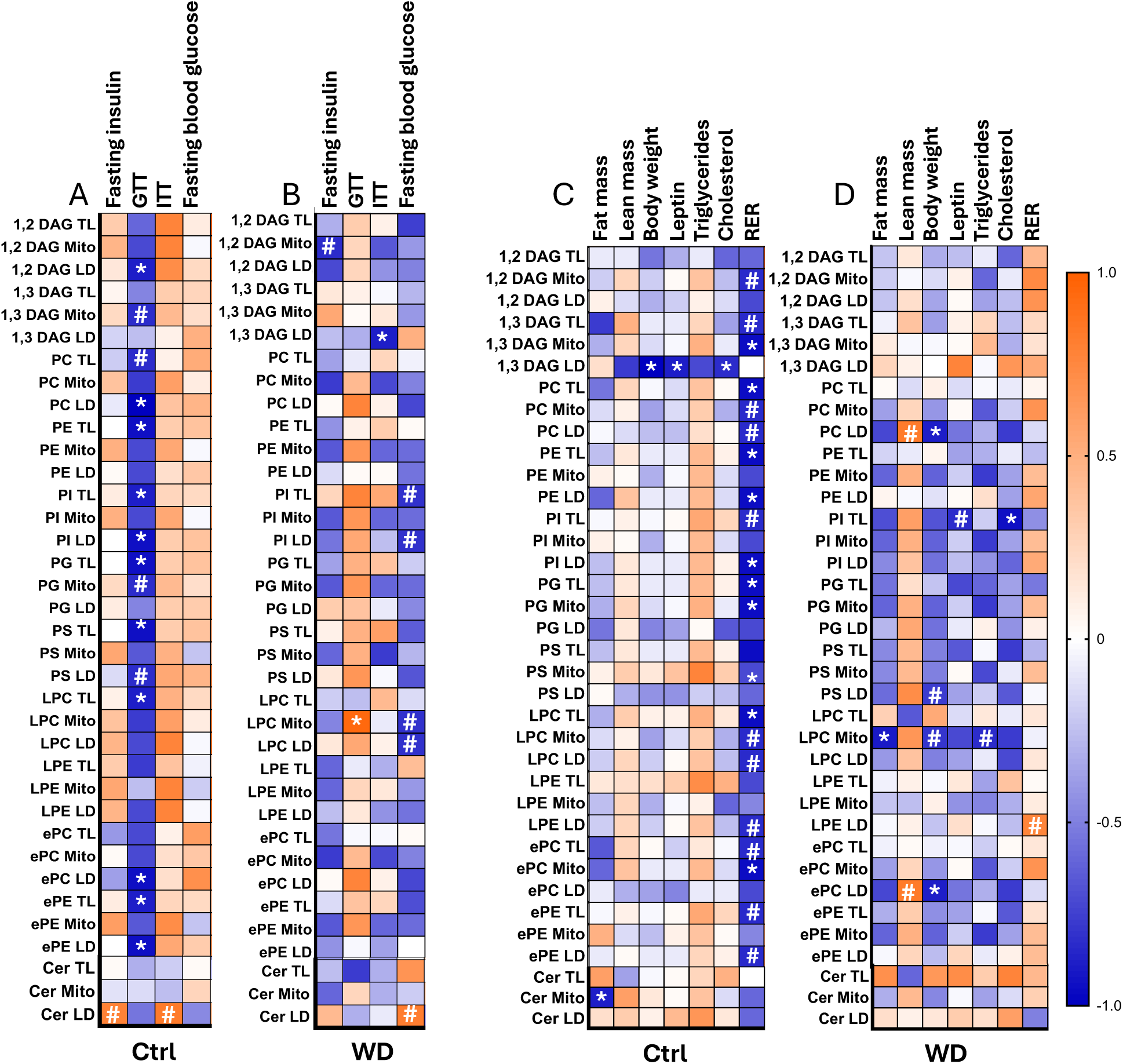
Correlations between lipid class abundance and metabolic or body composition parameters. (A–B) Spearman correlations between lipid classes and insulin resistance parameters in control (A) and WD-fed (B) mice. (C–D) Correlations between body composition measurements and lipid classes in control (C) and WD-fed (D) mice. Blue indicates a negative correlation and orange indicates a positive correlation, * p-value < 0.05, # p= 0.05-0.1. For all panels n=6 per group.

**Supplemental Table 1.**
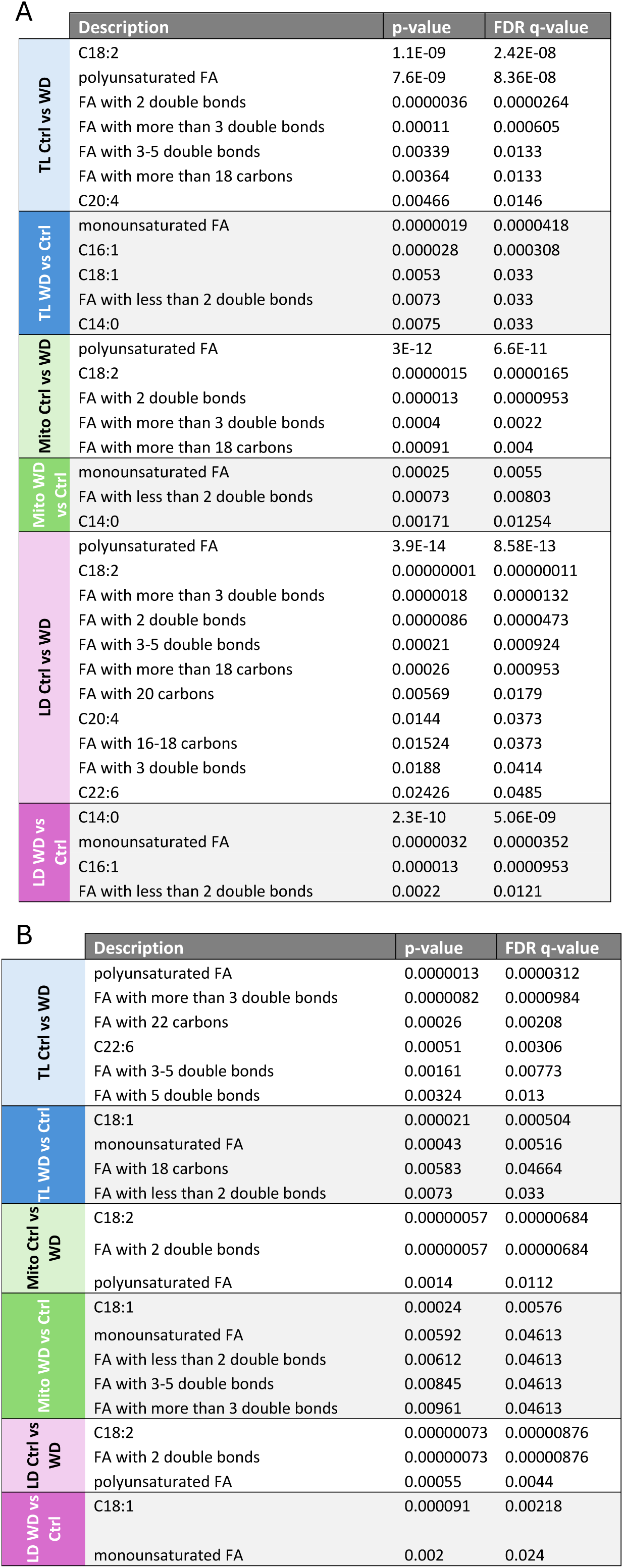
LION analysis. **A)** DAG and **B)** PL specific enrichment. Only categories with a FDR q-value < 0.05 were selected

## References

1. Aeberli, I., Hochuli, M., Gerber, P.A., Sze, L., Murer, S.B., Tappy, L., Spinas, G.A., and Berneis, K. (2013). Moderate amounts of fructose consumption impair insulin sensitivity in healthy young men: a randomized controlled trial. Diabetes Care 36, 150–156. 10.2337/dc12-0540.

2. Luukkonen, P.K., Sadevirta, S., Zhou, Y., Kayser, B., Ali, A., Ahonen, L., Lallukka, S., Pelloux, V., Gaggini, M., Jian, C., et al. (2018). Saturated Fat Is More Metabolically Harmful for the Human Liver Than Unsaturated Fat or Simple Sugars. Diabetes Care 41, 1732–1739. 10.2337/dc18-0071.

3. Nowotny, B., Zahiragic, L., Krog, D., Nowotny, P.J., Herder, C., Carstensen, M., Yoshimura, T., Szendroedi, J., Phielix, E., Schadewaldt, P., et al. (2013). Mechanisms underlying the onset of oral lipid-induced skeletal muscle insulin resistance in humans. Diabetes 62, 2240–2248. 10.2337/db12-1179.

4. Kien, C.L., Everingham, K.I., R, D.S., Fukagawa, N.K., and Muoio, D.M. (2011). Short-term effects of dietary fatty acids on muscle lipid composition and serum acylcarnitine profile in human subjects. Obesity (Silver Spring) 19, 305–311. 10.1038/oby.2010.135.

5. Kissebah, A.H., and Krakower, G.R. (1994). Regional adiposity and morbidity. Physiol Rev 74, 761–811. 10.1152/physrev.1994.74.4.761.

6. Coen, P.M., and Goodpaster, B.H. (2012). Role of intramyocelluar lipids in human health. Trends Endocrinol Metab 23, 391–398. 10.1016/j.tem.2012.05.009.

7. Jayasinghe, S.U., Tankeu, A.T., and Amati, F. (2019). Reassessing the Role of Diacylglycerols in Insulin Resistance. Trends Endocrinol Metab 30, 618–635. 10.1016/j.tem.2019.06.005.

8. Vance, D.E., and Vance, J.E. (2008). Biochemistry of lipids, lipoproteins and membranes, 5th Edition (Elsevier).

9. Penno, A., Hackenbroich, G., and Thiele, C. (2013). Phospholipids and lipid droplets. Biochim Biophys Acta 1831, 589–594. 10.1016/j.bbalip.2012.12.001.

10. Funai, K., Summers, S.A., and Rutter, J. (2020). Reign in the membrane: How common lipids govern mitochondrial function. Curr Opin Cell Biol 63, 162–173. 10.1016/j.ceb.2020.01.006.

11. Goni, F.M., and Alonso, A. (1999). Structure and functional properties of diacylglycerols in membranes. Prog Lipid Res 38, 1–48. 10.1016/s0163-7827(98)00021-6.

12. Eichmann, T.O., and Lass, A. (2015). DAG tales: the multiple faces of diacylglycerol--stereochemistry, metabolism, and signaling. Cell Mol Life Sci 72, 3931–3952. 10.1007/s00018-015-1982-3.

13. Perreault, L., Newsom, S.A., Strauss, A., Kerege, A., Kahn, D.E., Harrison, K.A., Snell-Bergeon, J.K., Nemkov, T., D’Alessandro, A., Jackman, M.R., et al. (2018). Intracellular localization of diacylglycerols and sphingolipids influences insulin sensitivity and mitochondrial function in human skeletal muscle. JCI Insight 3. 10.1172/jci.insight.96805.

14. Eum, J.Y., Lee, G.B., Yi, S.S., Kim, I.Y., Seong, J.K., and Moon, M.H. (2020). Lipid alterations in the skeletal muscle tissues of mice after weight regain by feeding a high-fat diet using nanoflow ultrahigh performance liquid chromatography-tandem mass spectrometry. J Chromatogr B Analyt Technol Biomed Life Sci 1141, 122022. 10.1016/j.jchromb.2020.122022.

15. Elshareif, N., Gornick, E., Gavini, C.K., Aubert, G., and Mansuy-Aubert, V. (2023). Comparison of western diet-induced obesity and streptozotocin mouse models: insights into energy balance, somatosensory dysfunction, and cardiac autonomic neuropathy. Front Physiol 14, 1238120. 10.3389/fphys.2023.1238120.

16. Gavini, C.K., Cook, T.M., Rademacher, D.J., and Mansuy-Aubert, V. (2020). Hypothalamic C2-domain protein involved in MC4R trafficking and control of energy balance. Metabolism 102, 153990. 10.1016/j.metabol.2019.153990.

17. Cook, T.M., Gavini, C.K., Jesse, J., Aubert, G., Gornick, E., Bonomo, R., Gautron, L., Layden, B.T., and Mansuy-Aubert, V. (2021). Vagal neuron expression of the microbiota-derived metabolite receptor, free fatty acid receptor (FFAR3), is necessary for normal feeding behavior. Mol Metab 54, 101350. 10.1016/j.molmet.2021.101350.

18. Arribat, Y., Broskey, N.T., Greggio, C., Boutant, M., Conde Alonso, S., Kulkarni, S.S., Lagarrigue, S., Carnero, E.A., Besson, C., Canto, C., and Amati, F. (2019). Distinct patterns of skeletal muscle mitochondria fusion, fission and mitophagy upon duration of exercise training. Acta Physiol (Oxf) 225, e13179. 10.1111/apha.13179.

19. Schindelin, J., Arganda-Carreras, I., Frise, E., Kaynig, V., Longair, M., Pietzsch, T., Preibisch, S., Rueden, C., Saalfeld, S., Schmid, B., et al. (2012). Fiji: an open-source platform for biological-image analysis. Nat Methods 9, 676–682. 10.1038/nmeth.2019.

20. Matyash, V., Liebisch, G., Kurzchalia, T.V., Shevchenko, A., and Schwudke, D. (2008). Lipid extraction by methyl-tert-butyl ether for high-throughput lipidomics. J Lipid Res 49, 1137–1146. 10.1194/jlr.D700041-JLR200.

21. Chassen, S.S., Zemski-Berry, K., Raymond-Whish, S., Driver, C., Hobbins, J.C., and Powell, T.L. (2022). Altered Cord Blood Lipid Concentrations Correlate with Birth Weight and Doppler Velocimetry of Fetal Vessels in Human Fetal Growth Restriction Pregnancies. Cells 11. 10.3390/cells11193110.

22. Harrison, K.A., and Bergman, B.C. (2019). HPLC-MS/MS Methods for Diacylglycerol and Sphingolipid Molecular Species in Skeletal Muscle. Methods Mol Biol 1978, 137–152. 10.1007/978-1-4939-9236-2_9.

23. Grepper, D., Tabasso, C., Zanou, N., Aguettaz, A.K.F., Castro-Sepulveda, M., Ziegler, D.V., Lagarrigue, S., Arribat, Y., Martinotti, A., Ebrahimi, A., et al. (2024). BCL2L13 at endoplasmic reticulum-mitochondria contact sites regulates calcium homeostasis to maintain skeletal muscle function. iScience 27, 110510. 10.1016/j.isci.2024.110510.

24. Pang, Z., Lu, Y., Zhou, G., Hui, F., Xu, L., Viau, C., Spigelman, A.F., MacDonald, P.E., Wishart, D.S., Li, S., and Xia, J. (2024). MetaboAnalyst 6.0: towards a unified platform for metabolomics data processing, analysis and interpretation. Nucleic Acids Res 52, W398–W406. 10.1093/nar/gkae253.

25. Molenaar, M.R., Jeucken, A., Wassenaar, T.A., van de Lest, C.H.A., Brouwers, J.F., and Helms, J.B. (2019). LION/web: a web-based ontology enrichment tool for lipidomic data analysis. Gigascience 8. 10.1093/gigascience/giz061.

26. Nie, Y., Gavin, T.P., and Kuang, S. (2015). Measurement of Resting Energy Metabolism in Mice Using Oxymax Open Circuit Indirect Calorimeter. Bio Protoc 5. 10.21769/bioprotoc.1602.

27. Yuping Yuan, S.M.S.H.H.S., and Shaun, P.J. (1996). The Bioactive Phospholipid, Lysophosphatidylcholine, Induces Cellular Effects via G-Protein-dependent Activation of Adenylyl Cyclase. The Journal of Biological Chemistry 271, 27090–27098.

28. Garcia-Jaramillo, M., Spooner, M.H., Lohr, C.V., Wong, C.P., Zhang, W., and Jump, D.B. (2019). Lipidomic and transcriptomic analysis of western diet-induced nonalcoholic steatohepatitis (NASH) in female Ldlr -/- mice. PLoS One 14, e0214387. 10.1371/journal.pone.0214387.

29. Busnelli, M., Manzini, S., Colombo, A., Franchi, E., Laaperi, M., Laaksonen, R., and Chiesa, G. (2024). Effect of diet and genotype on the lipidome of mice with altered lipoprotein metabolism. iScience 27, 111051. 10.1016/j.isci.2024.111051.

30. Heintz, M.M., Kumar, R., Maner-Smith, K.M., Ortlund, E.A., and Baldwin, W.S. (2022). Age- and Diet-Dependent Changes in Hepatic Lipidomic Profiles of Phospholipids in Male Mice: Age Acceleration in Cyp2b-Null Mice. J Lipids 2022, 7122738. 10.1155/2022/7122738.

31. Chen, Z., Ho, I.L., Soeung, M., Yen, E.Y., Liu, J., Yan, L., Rose, J.L., Srinivasan, S., Jiang, S., Edward Chang, Q., et al. (2023). Ether phospholipids are required for mitochondrial reactive oxygen species homeostasis. Nat Commun 14, 2194. 10.1038/s41467-023-37924-9.

32. Goodpaster, B.H., Thaete, F.L., Simoneau, J.A., and Kelley, D.E. (1997). Subcutaneous abdominal fat and thigh muscle composition predict insulin sensitivity independently of visceral fat. Diabetes 46, 1579–1585. 10.2337/diacare.46.10.1579.

33. Phillips, D.I., Caddy, S., Ilic, V., Fielding, B.A., Frayn, K.N., Borthwick, A.C., and Taylor, R. (1996). Intramuscular triglyceride and muscle insulin sensitivity: evidence for a relationship in nondiabetic subjects. Metabolism 45, 947–950. 10.1016/s0026-0495(96)90260-7.

34. Renier, T.J., Paetz, O.R., Paal, M.C., Long, A.B., Brown, M.R., Vuong, S.H., Perumal, S.K., Kharbanda, K.K., and Listenberger, L.L. (2023). Changing the phospholipid composition of lipid droplets alters localization of select lipid droplet proteins. MicroPubl Biol 2023. 10.17912/micropub.biology.000960.

35. Wolk, M., and Fedorova, M. (2024). The lipid droplet lipidome. FEBS Lett 598, 1215–1225. 10.1002/1873-3468.14874.

36. Goodpaster, B.H., Wolfe, R.R., and Kelley, D.E. (2002). Effects of obesity on substrate utilization during exercise. Obes Res 10, 575–584. 10.1038/oby.2002.78.

37. Benador, I.Y., Veliova, M., Liesa, M., and Shirihai, O.S. (2019). Mitochondria Bound to Lipid Droplets: Where Mitochondrial Dynamics Regulate Lipid Storage and Utilization. Cell Metab 29, 827–835. 10.1016/j.cmet.2019.02.011.

38. Lyu, K., Zhang, Y., Zhang, D., Kahn, M., Ter Horst, K.W., Rodrigues, M.R.S., Gaspar, R.C., Hirabara, S.M., Luukkonen, P.K., Lee, S., et al. (2020). A Membrane-Bound Diacylglycerol Species Induces PKCɛ-Mediated Hepatic Insulin Resistance. Cell Metab 32, 654–664 e655. 10.1016/j.cmet.2020.08.001.

39. Petersen, M.C., Yoshino, M., Smith, G.I., Gaspar, R.C., Kahn, M., Samovski, D., Shulman, G.I., and Klein, S. (2024). Effect of Weight Loss on Skeletal Muscle Bioactive Lipids in People With Obesity and Type 2 Diabetes. Diabetes 73, 2055–2064. 10.2337/db24-0083.

40. Saito, S., Hernandez-Ono, A., and Ginsberg, H.N. (2007). Dietary 1,3-diacylglycerol protects against diet-induced obesity and insulin resistance. Metabolism 56, 1566–1575. 10.1016/j.metabol.2007.06.024.

41. Bergman, B.C., Perreault, L., Strauss, A., Bacon, S., Kerege, A., Harrison, K., Brozinick, J.T., Hunerdosse, D.M., Playdon, M.C., Holmes, W., et al. (2018). Intramuscular triglyceride synthesis: importance in muscle lipid partitioning in humans. Am J Physiol Endocrinol Metab 314, E152–E164. 10.1152/ajpendo.00142.2017.

42. Gaspar, R.C., Lyu, K., Hubbard, B.T., Leitner, B.P., Luukkonen, P.K., Hirabara, S.M., Sakuma, I., Nasiri, A., Zhang, D., Kahn, M., et al. (2023). Distinct subcellular localisation of intramyocellular lipids and reduced PKCepsilon/PKCtheta activity preserve muscle insulin sensitivity in exercise-trained mice. Diabetologia 66, 567–578. 10.1007/s00125-022-05838-8.

43. Amati, F., Dube, J.J., Alvarez-Carnero, E., Edreira, M.M., Chomentowski, P., Coen, P.M., Switzer, G.E., Bickel, P.E., Stefanovic-Racic, M., Toledo, F.G., and Goodpaster, B.H. (2011). Skeletal muscle triglycerides, diacylglycerols, and ceramides in insulin resistance: another paradox in endurance-trained athletes? Diabetes 60, 2588–2597. 10.2337/db10-1221.

44. Goto-Inoue, N., Yamada, K., Inagaki, A., Furuichi, Y., Ogino, S., Manabe, Y., Setou, M., and Fujii, N.L. (2013). Lipidomics analysis revealed the phospholipid compositional changes in muscle by chronic exercise and high-fat diet. Sci Rep 3, 3267. 10.1038/srep03267.

45. Matsuzaka, T., Shimano, H., Yahagi, N., Kato, T., Atsumi, A., Yamamoto, T., Inoue, N., Ishikawa, M., Okada, S., Ishigaki, N., et al. (2007). Crucial role of a long-chain fatty acid elongase, Elovl6, in obesity-induced insulin resistance. Nat Med 13, 1193–1202. 10.1038/nm1662.

46. Rahman, S.M., Dobrzyn, A., Dobrzyn, P., Lee, S.H., Miyazaki, M., and Ntambi, J.M. (2003). Stearoyl-CoA desaturase 1 deficiency elevates insulin-signaling components and down-regulates protein-tyrosine phosphatase 1B in muscle. Proc Natl Acad Sci U S A 100, 11110–11115. 10.1073/pnas.1934571100.

47. Vessby, B., Gustafsson, I.B., Tengblad, S., Boberg, M., and Andersson, A. (2002). Desaturation and elongation of fatty acids and insulin action. Annals of the New York Academy of Sciences 967, 183–195. 10.1111/j.1749-6632.2002.tb04275.x.

48. Joshi, A., Richard, T.H., and Gohil, V.M. (2023). Mitochondrial phospholipid metabolism in health and disease. J Cell Sci 136. 10.1242/jcs.260857.

49. Wong, S., Bertram, K.R., Balasubramaniam, S.S., Raghuram, N., Knight, T., Maschek, J.A., Cox, J.E., and Hughes, A.L. (2025). Alterations in lipid saturation trigger remodeling of the outer mitochondrial membrane. Mol Biol Cell 36, ar129. 10.1091/mbc.E25-01-0033.

50. Hoeks, J., Wilde, J., Hulshof, M.F., Berg, S.A., Schaart, G., Dijk, K.W., Smit, E., and Mariman, E.C. (2011). High fat diet-induced changes in mouse muscle mitochondrial phospholipids do not impair mitochondrial respiration despite insulin resistance. PLoS One 6, e27274. 10.1371/journal.pone.0027274.

